# Multimodal dynamics during naturalistic learning are associated with subsequent performance

**DOI:** 10.64898/2026.11.23.768566

**Authors:** Ziqiang Tian, An Long, Yaqiong Xiao

## Abstract

Learning from continuous material involves dynamic coordination of visual sampling, autonomic arousal, and neural activity, yet how these signals jointly organize during successful learning remains unclear. Here, we analysed synchronized eye-tracking, pupillometry, and EEG recordings from 86 participants from the Brain, Body, and Behavior Dataset. Using hidden Markov modelling, we identified recurrent multimodal states after accounting for stable recording-level differences and stimulus-shared responses. Three reproducible states with distinct psychophysiological profiles emerged. Poorer subsequent quiz performance was associated with greater relative expression of a state characterized by longer fixations, reduced fixation and saccade rates, lower gaze entropy, elevated pupil level and variability, and distinct EEG spectral entropy and frontoparietal theta coherence. In contrast, better performance was associated with greater expression of a state characterized primarily by more frequent and larger phasic pupil dilations. Quiz performance was also positively associated with transitions from this phasic-pupil state to the dominant background state, whereas relative expression of the dominant state itself showed no robust association with performance. The transition association did not survive correction in common-record window-length sensitivity analyses. These findings indicate that learning during naturalistic viewing is associated with the dynamic coordination of multiple ocular, pupillary, and neural signals, with relative state balance and directed transition probabilities each associated with subsequent quiz performance.

## 1 Introduction

Learning from naturalistic experience depends not only on whether learners are generally attentive, but also on how attention and related physiological processes fluctuate over time. When watching a cooking video, for example, we may feel focused yet later realize that we remember some steps clearly while having little memory for others. Such an experience raises a broader question: what aspects of moment-to-moment processing distinguish simply watching from learning? Behavioural and neural fluctuations accompany changes in sustained attention during continuous performance tasks (Esterman et al., 2013), and attention modulates the reliability of neural responses to naturalistic narratives (Ki et al., 2016). Neural engagement during educational videos predicts subsequent retention (Cohen et al., 2018), synchronized eye movements are associated with test performance (Madsen et al., 2021), and electroencephalography (EEG)-derived attentional state can predict learning outcomes (Madsen & Parra, 2025). These findings indicate that moment-to-moment variations in attention and physiology are relevant to learning, but leave open how visual sampling, arousal-related pupil dynamics and neural activity are jointly organized within a learner over time. Understanding this joint organization may reveal aspects of learning-related processing that are not captured by measures of attention or engagement considered in isolation. We therefore asked whether recurring multimodal configurations, their relative expression over time, and transitions between them are associated with subsequent learning.

Naturalistic learning provides a particularly informative context for examining this question because visual sampling, pupil dynamics, and neural activity fluctuate continuously and concurrently. Importantly, these signals do not map uniquely onto separate cognitive processes. Eye movements reflect how learners sample visual information, but their cognitive interpretation depends on the material and task. A review of video-learning research highlighted the difficulty of disentangling eye-tracking metrics from the cognitive processes they are often assumed to represent (Deng & Gao, 2023). Similarly, during movie viewing, neural oscillations covary with both subjective and pupillary measures of arousal (Camenzind et al., 2026). During informative videos, EEG, gaze, pupil size, and heart rate exhibit intersubject synchrony that varies with attention and predicts subsequent recall (Madsen & Parra, 2022), while multimodal eye tracking and functional imaging have linked instructional conditions to visual attention, neural synchrony, and post-test performance (Gu et al., 2024). These findings establish meaningful relationships among visual, autonomic, and neural signals during naturalistic viewing. Yet they do not reveal whether these signals form recurring configurations within individuals as learning unfolds. The critical unresolved question is therefore not simply whether particular signals relate to learning, but whether their joint temporal organization provides a useful description of learning-related dynamics.

A latent-state approach offers a way to characterize such organization without assigning fixed cognitive labels to individual signals. Rather than treating multimodal measures as independent time series, this approach asks whether observations repeatedly occupy a limited set of configurations and whether the occupancy and transitions of these configurations follow structured temporal patterns. It distinguishes the relative expression of different configurations from the sequence in which configurations occur. Switching dynamical models have identified hidden neural states whose recruitment and transitions are associated with cognitive performance (Taghia et al., 2018). Movie viewing has also been shown to elicit reproducible brain-state dynamics aligned with aspects of the unfolding stimulus (Meer et al., 2020), while neural trajectories in shared low-dimensional spaces vary across cognitive and attentional conditions (Song et al., 2023). Related work on narrative event segmentation and environmental-state updating further illustrates the behavioural relevance of temporal organization, although these constructs differ from recurrent physiological states (Baldassano et al., 2017; Li et al., 2026). Hidden Markov models (HMMs) provide a framework for characterizing such recurring configurations and their transitions in continuous recordings without requiring predefined trial boundaries (Versteijnen & Heimel, 2026). These advances leave open whether eye-movement, pupil, and EEG fluctuations jointly organize into recurrent multimodal configurations during educational viewing, and whether differences in their relative expression and directed transitions are associated with subsequent learning.

Addressing this question requires synchronized multimodal measurements obtained during the same viewing episodes, together with behavioural measures of learning. The Brain, Body, and Behavior Dataset (BBBD) provides aligned ocular, autonomic, and neural recordings collected during educational videos and accompanying measures of learning (Madsen et al., 2026). The common stimulus timeline also makes it possible to distinguish activity shared across viewers from participant-specific variation. Shared responses during naturalistic viewing can be behaviourally meaningful: stronger intersubject correlations during movie viewing have been associated with successful episodic encoding (Hasson et al., 2008), and physiological synchrony during informative videos varies with attention and predicts recall (Madsen & Parra, 2022). However, a multimodal configuration occurring at the same scene across viewers may partly reflect stimulus-driven processing that is common across viewers rather than individual learning-related variation. We therefore examined participant-specific multimodal fluctuations after accounting for stable recording-level differences and shared video-related responses, asking how residual multimodal activity organizes over time and whether this organization is associated with subsequent quiz performance.

Here, we analysed simultaneous eye-movement, pupil, and EEG recordings from 86 participants across Experiments 2 to 4 of the BBBD, comprising 304 participant-by-video recordings. Our objective was to characterize recurrent multimodal configurations during educational viewing and examine whether their state occupancy and temporal transitions were associated with subsequent quiz performance. Specifically, we addressed three questions: (1) whether reproducible multimodal states with interpretable feature profiles could be identified; (2) whether individual differences in the relative expression of these states were associated with quiz performance; and (3) whether directed transitions between states were associated with performance. We expected that learning-related variation would be reflected in the joint organization of visual sampling, pupil dynamics, and neural activity. We treated these as exploratory, non-directional expectations rather than prespecified predictions about particular states or paths. Using an unsupervised HMM framework, we characterized the latent multimodal organization of eye movements, pupil dynamics, and neural activity during movie viewing and examined its associations with learning outcomes. By moving beyond isolated measures of attention or arousal, this approach captures how these signals jointly organize over time during naturalistic learning.

## 2 Methods

### 2.1 Dataset and experimental design

#### 2.1.1 Dataset and participants

We analysed data from the BBBD, which contains synchronized multimodal recordings acquired during naturalistic video viewing (Madsen et al., 2026). The present analysis included 86 participants from Experiments 2–4 (Experiment 2: *n* = 27; Experiment 3: *n* = 25; Experiment 4: *n* = 34). These participants contributed 304 valid participant-by-video recordings across 14 experiment-specific video IDs representing 11 unique video contents. Figure 1 summarizes the experimental design and analysis workflow.

**Figure 1.**
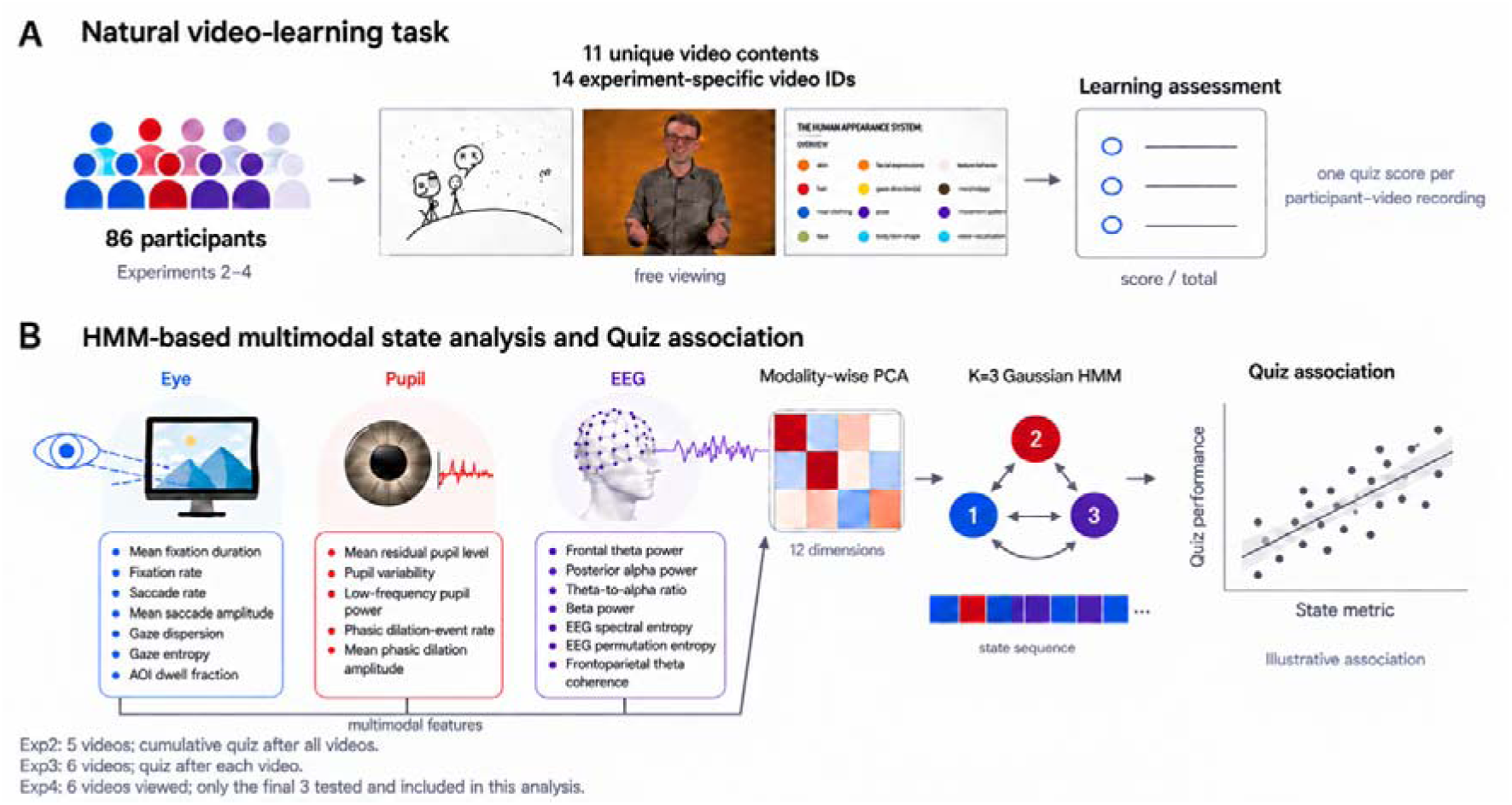
Experimental design and analysis workflow. (A) Experiment-specific viewing and quiz schedules for the analysed sample. Eleven unique video contents corresponded to 14 experiment-specific video IDs (5 + 6 + 3). (B) Eye-movement, pupil and EEG features from 30-s windows were reduced to 12 modality-specific principal components and entered into a hidden Markov model (HMM) to identify recurring multimodal temporal states, illustrated schematically as three states. The resulting features were then related to quiz performance. State transitions and the positive association depicted on the right are schematic and do not represent prespecified state-specific predictions or empirical results. EEG, electroencephalography; PCA, principal component analysis; HMM, hidden Markov model.

#### 2.1.2 Experimental paradigm and behavioural measure

Participants viewed educational videos covering astronomy, biology, physics, and science and technology, with unconstrained gaze. The analysis contained five video IDs from Experiment 2, six from Experiment 3 and three from Experiment 4. The three Experiment 4 contents also occurred in Experiments 2 or 3, yielding 11 unique contents across 14 experiment–video combinations. Across these 14 combinations, processed stimulus durations averaged 4.83 min (range: 2.38–6.48 min).

Quiz procedures differed across experiments. In Experiment 2, participants viewed five videos without advance notice of testing and subsequently completed a cumulative quiz. In Experiment 3, participants were informed that they would be tested and completed a quiz after each of six videos. In Experiment 4, participants viewed six videos without advance notice of testing; only the final three were followed by quizzes, and these three videos provided the recordings and scores analysed here. Video-specific quiz scores were obtained from the released questionnaire tables, including the per-video memory subscores of the cumulative quiz in Experiment 2. Each score was normalized by its corresponding maximum possible score, while retaining the scoring procedures specified in the dataset, including fractional item scores where applicable.

### 2.2 Multimodal data preprocessing and feature extraction

#### 2.2.1 Eye-tracking preprocessing

Gaze data were obtained from the preprocessed BBBD eye-tracking derivatives. Signals were recorded at 500 Hz with an EyeLink 1000 system, aligned to the video timeline and resampled to 128 Hz. Fixation and saccade events were obtained from the EyeLink annotations distributed with the dataset.

Samples marked as blinks, tracking artefacts, or unreliable measurements in the BBBD masks were excluded from window-level calculations. Extended gaps were not additionally interpolated. Windows were retained when at least 60% of gaze samples were valid and at least three fixations were available. Seven eye-tracking features were extracted from each window: mean fixation duration, fixation rate, saccade rate, mean saccade amplitude, gaze dispersion, gaze entropy, and area of interest (AOI) dwell fraction.

#### 2.2.2 Pupillometry preprocessing

Pupil analyses used the preprocessed pupil-size derivatives and the accompanying blink and interpolation annotations. In the original BBBD pipeline, pupil artefacts were identified as observations exceeding four times the interquartile range of a detrended pupil signal obtained using a 200-ms median filter. Blinks, detected artefacts and the surrounding 100-ms intervals were corrected by linear interpolation. The released processing code additionally accounts for changes in head-to-tracker distance when deriving the pupil signal.

In the present analysis, samples identified by the BBBD interpolation masks were first treated as missing. Only internal gaps of 500 ms or less were re-interpolated, whereas longer gaps remained missing. The cleaned pupil signal was then aggregated into 100-ms bins using the median, yielding a 10-Hz time series. Pupil quality control required at least 80% finite samples in the 10-Hz residual signal within each window, after short-gap interpolation and luminance adjustment.

Because video viewing did not include a common prestimulus baseline, no event-locked baseline correction was applied. To reduce variation attributable to visual stimulation and slow temporal trends, pupil size was residualized using ridge regression with video luminance at lags of 0.5–3.0 s, concurrent image contrast, scene changes and linear time trends as predictors. Luminance adjustment was included because luminance is a major time-varying determinant of pupil size in naturalistic stimuli (Fink et al., 2023; Kret & Sjak-Shie, 2019).

Five window-level pupil features were extracted: mean residual pupil level, pupil variability, low-frequency pupil power, phasic dilation-event rate, and mean phasic dilation amplitude. Pupil variability was quantified using the median absolute deviation, scaled by 1.4826. Low-frequency pupil power was calculated over 0.05–0.20 Hz using Welch spectral estimation and was log-transformed. Phasic dilation events were identified from robustly thresholded positive peaks in the smoothed pupil derivative, with a minimum separation of 1 s. Event amplitude was defined as the maximum residual pupil value during the 1.5 s following an event minus the median value during the preceding 0.5 s.

#### 2.2.3 EEG data preprocessing

EEG analyses used the preprocessed continuous EEG derivatives released by BBBD. EEG was originally recorded at 2,048 Hz using a 64-channel BioSemi ActiveTwo system. According to the BIDS sidecars accompanying the files used in the present analysis, the signals were high-pass filtered at 0.3 Hz and notch-filtered at 60 Hz using fifth-order Butterworth filters. The BBBD pipeline applied robust principal component analysis to suppress sparse non-physiological artefacts, removed ocular activity through least-squares regression of the EOG channels, interpolated outlying samples and their surrounding 40-ms intervals using neighbouring electrodes, low-pass filtered the data at 64 Hz and downsampled the cleaned signals to 128 Hz. No additional independent component analysis (ICA) or manual artefact correction was applied in the present study. EEG features were extracted from ongoing activity within each analysis window using spectral and complexity measures.

Power spectral density was estimated within each analysis window using Welch’s method with 4-s Hann segments and 2-s overlap (Welch, 1967). Frontal theta power was calculated by integrating 4–8 Hz power over frontal and frontocentral electrodes, whereas posterior alpha power was calculated over 8–12 Hz using parietal and occipital electrodes. Band-power values were log-transformed.

The remaining EEG features comprised the theta-to-alpha ratio (log-transformed), beta power (global power over 13–30 Hz), EEG spectral entropy (normalized), EEG permutation entropy and frontoparietal theta coherence. Frequencies between 15 and 17 Hz were excluded from beta-power and EEG spectral-entropy calculations because BBBD documents a non-physiological 16-Hz electrical component in part of the dataset. Frontoparietal theta coherence was calculated as the mean magnitude-squared coherence between regional frontal and posterior signals over 4–8 Hz.

#### 2.2.4 Temporal segmentation and feature extraction

The synchronized eye-tracking, pupillometry, and EEG recordings were segmented into 30-s overlapping windows advanced in 10-s increments. This window length was chosen to provide stable estimation of eye-movement rates, low-frequency pupil fluctuations, and EEG spectral features while retaining temporal resolution across each video.

Windows that did not meet the modality-specific quality-control criteria were excluded from subsequent analyses. Missing windows were retained as gaps, such that the remaining valid windows formed separate continuous sequences within each participant-by-video recording. After multimodal quality control, 5,302 overlapping windows were retained, corresponding to 17.55 h of non-overlapping temporal coverage.

### 2.3 Hidden Markov modelling of multimodal states

#### 2.3.1 Construction of the HMM feature space

For HMM estimation, continuous sequences containing fewer than three valid windows were excluded. The remaining sequences were entered into the HMM while preserving their temporal order; state transitions were modelled only between consecutive valid windows within the same continuous sequence. Before model fitting, each feature was mean centred separately within each participant and video to remove stable participant- and video-level differences while preserving within-recording fluctuations.

To further remove responses shared across viewers, which may reflect the timing and content of the video rather than participant-specific fluctuations, a leave-one-participant-out group time course was calculated for every experiment, video, and window following the logic of inter-subject response modelling in naturalistic paradigms (Nastase et al., 2019). For each feature, a single regression slope was estimated by pooling the available centred observations and their corresponding leave-one-participant-out group values across participants and videos; the scaled shared response was subtracted from each observation. Residual features were then standardized within each participant and video using the median and median absolute deviation (MAD), and restricted to the range -8 to 8 to limit the influence of extreme values. The resulting features represented participant-specific within-recording fluctuations after removing stable participant/video differences and responses shared across viewers.

The feature set comprised seven eye-movement features, five pupil features, and seven EEG features. Dimensionality reduction was performed separately within each modality to prevent the modality with the largest number of original variables from disproportionately influencing state estimation. Within each modality, standardized features were entered into principal component analysis (PCA), and the smallest number of components explaining at least 80% of the modality-specific variance was retained, subject to a maximum of four components. Four components were retained for each modality, yielding a 12-dimensional representation comprising four eye-tracking, four pupil, and four EEG components.

For participant-held-out validation, shared video time courses, feature-specific regression slopes and PCA transformations were estimated using training participants only and applied to held-out participants. Median/MAD standardization was performed separately within each participant-by-video recording, including held-out recordings; it did not use quiz outcomes.

#### 2.3.2 Hidden Markov model estimation and state-number selection

A first-order Gaussian HMM with diagonal covariance was fitted to the 12-dimensional multimodal representation (Taghia et al., 2018; Vidaurre et al., 2016). The model provided posterior state probabilities for each window and the corresponding most-probable state sequence. State labels were ordered by overall posterior occupancy, with State 1 denoting the most frequently expressed state. These labels did not imply an a priori cognitive ranking; state interpretation was based on the features characterizing each state.

The number of states was selected in two stages because HMM solutions can vary across model specifications and random initializations (Alonso & Vidaurre, 2023; Nielsen et al., 2018). Models with two to eight states were fitted from 20 random initializations. Solutions were excluded if they failed to converge, contained a state with occupancy below 5%, or had a median adjusted Rand index below 0.75 across initializations. These criteria were prespecified as study-specific safeguards against unstable or poorly represented solutions. Among admissible models, the Bayesian information criterion (BIC) was used as the primary complexity-penalized fit measure (Schwarz, 1978). Akaike information criterion (AIC; Akaike, 1974) and participant-held-out log likelihood were used as secondary measures of model fit and participant-level generalization, respectively (Nielsen et al., 2018).

#### 2.3.3 Characterization of the multimodal states

States were characterized using the original 19 features rather than the PCA scores. For each participant and state, a posterior-weighted feature mean was calculated by weighting every window by the probability of belonging to that state. Estimates were retained when the accumulated posterior weight was equivalent to at least three windows.

Differences in feature values across states were examined using Friedman tests, followed by two-sided paired-sample *t*-tests for pairwise comparisons. The Benjamini–Hochberg procedure controlled the false discovery rate (FDR) separately across the 19 omnibus tests and the 57 pairwise comparisons (Benjamini & Hochberg, 1995). We summarize eligible participant-level state estimates using means and sample standard deviations and report both unadjusted p values and FDR-adjusted q values for omnibus and pairwise comparisons.

#### 2.3.4 State dynamics

The temporal expression and organization of the identified states were characterized using state occupancy, mean dwell time, and transition probabilities. State occupancy was defined as the proportion of windows assigned to each state in the most-probable state sequence. Mean dwell time was calculated from consecutive runs of the same state label using the 10-s step (Larsen et al., 2026; Taghia et al., 2018). Transition probabilities were estimated from consecutive state labels. Self-transitions were excluded when analysing switching destinations. For each directed between-state transition, participants with no observed switch from the source state were assigned a value of 0; otherwise, transition probability was defined as the proportion of observed switches from the source state that reached the specified destination state.

Differences in state occupancy and mean dwell time across the three states were examined using Friedman tests, followed by two-sided paired-sample *t*-tests. Complete cases were used for each omnibus test, whereas each paired comparison used participants with both required values. Benjamini–Hochberg FDR correction was applied separately across the three pairwise comparisons for each metric. In an additional exploratory analysis, the six zero-inclusive between-state transition probabilities were compared using a Friedman test followed by all 15 paired-sample *t*-tests, with FDR correction across those 15 comparisons. Omnibus tests are reported with unadjusted *p* values; pairwise comparisons are reported with *t* statistics, degrees of freedom, unadjusted *p* values and FDR-adjusted *q* values. These transition comparisons describe differences among observed destination probabilities, including zeros assigned when no source-state switch occurred; they do not test differences in switching frequency.

### 2.4 Associations with quiz performance

#### 2.4.1 Relative state balance and quiz performance

We tested whether individual differences in the relative posterior occupancy of each state were associated with quiz performance. State occupancies were normalized across the three states and therefore summed to one, making them compositional and unsuitable for inclusion as independent predictors. To account for this constraint, we defined relative state balance for each target state as a log-ratio of its posterior occupancy to the geometric mean of the posterior occupancies of the other two states (Greenacre, 2021; Susin et al., 2020):

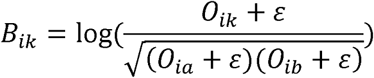

where O denotes the mean posterior state probability within a participant-by-video recording (posterior occupancy), *k* denotes the target state, and *a* and *b* denote the two non-target states. A small constant (10^-6^) was added to occupancies before calculating the log ratio to accommodate zero values. Positive values indicate relatively greater occupancy of the target state, whereas negative values indicate relatively lower occupancy relative to the other states. The resulting state-balance measures were averaged across videos for each participant and standardized before analysis.

For each target state, we fitted a participant-clustered fractional-logistic generalized estimating equation (GEE) model, with participant-by-video quiz proportion as the outcome. Models included standardized between-participant relative state balance, age, sex, and experiment-specific video identity (14 levels) as predictors. Relative state balance was entered only as a between-participant predictor because it was averaged across videos within each participant. Models used a binomial mean-variance family with a logit link, quiz-item count as observation weights, an independence working correlation structure, and robust sandwich standard errors to account for repeated video observations within participants. Coefficients represent the change in the log-odds of quiz performance associated with a 1-*SD* increase in state balance. Participants with an absolute standardized value greater than 3 for either mean direct occupancy or mean state balance were excluded separately for each target-state analysis; predictors were then recalculated in the retained sample.

#### 2.4.2 Directed state transitions and quiz performance

We next examined whether directed transitions between states were associated with quiz performance. Whereas state occupancy captures the relative expression of states over the viewing period, directed transitions characterize how one state followed another.

For each participant, quiz performance was calculated as the total points earned across videos divided by the total number of quiz items contributing to the participant’s outcome. For each directed transition (a → b), where a ≠ b, we counted switches between adjacent windows across all valid sequences, excluding transitions across sequence boundaries. Transition probability was defined as the proportion of observed switches out of state a that were directed to state b. Participants with no observed switch out of state a were assigned a value of zero. Thus, a zero value could reflect either no observed switching from the source state or no switching to the specified destination among observed switches, and therefore should not be interpreted as a direct measure of switching frequency.

A separate weighted least-squares regression model was fitted for each of the six directed transitions (1→2, 1→3, 2→1, 2→3, 3→1, and 3→2). Each model included the standardized transition probability and age as continuous predictors, with sex and experiment as categorical covariates. Quiz performance was weighted by the total number of quiz items contributing to each participant’s outcome. Models included participants with complete outcome, weight, age, sex, and transition data. HC3 heteroskedasticity-robust standard errors were used to construct 95% confidence intervals and two-sided tests. The transition coefficient represents the adjusted difference in quiz performance associated with a 1-*SD* increase in the transition probability.

#### 2.4.3 Multiple-comparison correction and statistical reporting

Benjamini–Hochberg FDR correction was applied separately to the three state-balance GEE models and the six directed-transition models. Statistical significance was assessed at *q* < .05 within each test family. Throughout, *p* denotes the two-sided model-test *p* value before multiplicity correction, and *q* denotes the corresponding Benjamini–Hochberg FDR-adjusted value. Both *p* and *q* values were derived from the covariate-adjusted models; thus, an uncorrected *p* value does not represent an unadjusted regression model. We report regression coefficients, two-sided p values, FDR-adjusted q values, and robust 95% confidence intervals (Benjamini & Hochberg, 1995).

### 2.5 Window-length sensitivity analysis

To assess whether the primary findings were sensitive to the choice of window length, we repeated the analysis using 20-s windows with the same 10-s step. To ensure that differences between the 20-s and 30-s solutions were not attributable to differences in the recordings included, the sensitivity analysis was restricted to the 279 participant-by-video recordings from 81 participants that yielded valid sequences at both window lengths. Within this common-recording sample, the full analysis pipeline was repeated separately for the 30-s and 20-s windows using the same feature definitions, quality-control procedures, centering, shared-response removal, and modality-specific PCA retention rule. The resulting representations contained 12 components for the 30-s windows and 11 components for the 20-s windows. Three-state diagonal-covariance Gaussian HMMs were fitted separately at each window length using 20 random initializations. The common-record 30-s solution was used as the reference for comparison with the 20-s sensitivity solution; it was not treated as a replacement for the primary full-sample 30-s analysis.

The three states in the common-record 30-s solution were first aligned with the corresponding states in the primary full-sample 30-s solution. The 20-s states were then aligned with the common-record 30-s states. State matching was based solely on the 19-feature state profiles, without reference to quiz performance or other outcome measures, using one-to-one assignment that maximized the summed Pearson correlations between matched profiles. Similarity between matched state profiles was quantified using Pearson correlations across the 19 features. We similarly compared the complete 3 × 3 row-normalized transition matrices, including self-transitions, between the common-record 30-s and 20-s solutions using Pearson correlation across all nine corresponding entries. Pearson correlation coefficients and their nominal two-sided *p* values are reported as descriptive measures of correspondence. Because the feature and matrix entries are not independent and state matching was performed to maximize correspondence, these *p* values should not be interpreted as formal inferential tests of robustness.

State-balance associations with quiz performance were re-estimated for the 20-s sensitivity solution using the same participant-clustered fractional-logistic GEE specification as in the primary state-balance analysis. For direct comparison, the same GEE analyses were also re-estimated for the common-record 30-s solution. FDR correction was applied across the three state-balance tests separately for each window length. Target-state exclusions were applied separately within each analysis, so the final GEE samples could differ despite the common starting set of participant-by-video recordings.

As an additional exploratory sensitivity analysis, the six directed transition–quiz associations were re-estimated for both the 20-s and common-record 30-s solutions using the same weighted least-squares models, covariates, quiz-item weights, and HC3 standard errors as in the primary transition–quiz analysis. FDR correction was applied across the six transition tests separately for each window length. These analyses used the common-recording samples and the same zero-inclusive definition of transition probability; no state-balance-specific exclusions were applied to the transition analyses.

## 3 Results

### 3.1 A stable three-state solution with distinct multimodal profiles

Only the two- and three-state models met the prespecified admissibility criteria. Compared with the two-state model, the three-state model showed a lower BIC (182,671.7 versus 183,967.6) and a higher participant-held-out log likelihood per window (-17.192 versus -17.316). We therefore retained the three-state solution. Models with four or more states were excluded because at least one state was unstable or infrequently expressed.

Three states showed distinct multimodal profiles (Figure 2). State 1 was the dominant configuration, with most feature values remaining close to the within-recording reference. State 2 was characterized by longer fixations, lower fixation and saccade rates, lower gaze entropy, higher pupil level and variability, higher posterior alpha power and frontoparietal theta coherence, and lower theta-to-alpha ratio and EEG spectral entropy. State 3 was distinguished primarily by more frequent and larger phasic pupil dilations, together with elevated pupil level and variability. Detailed feature estimates and statistical comparisons are provided in Supplementary Table S1.

**Figure 2.**
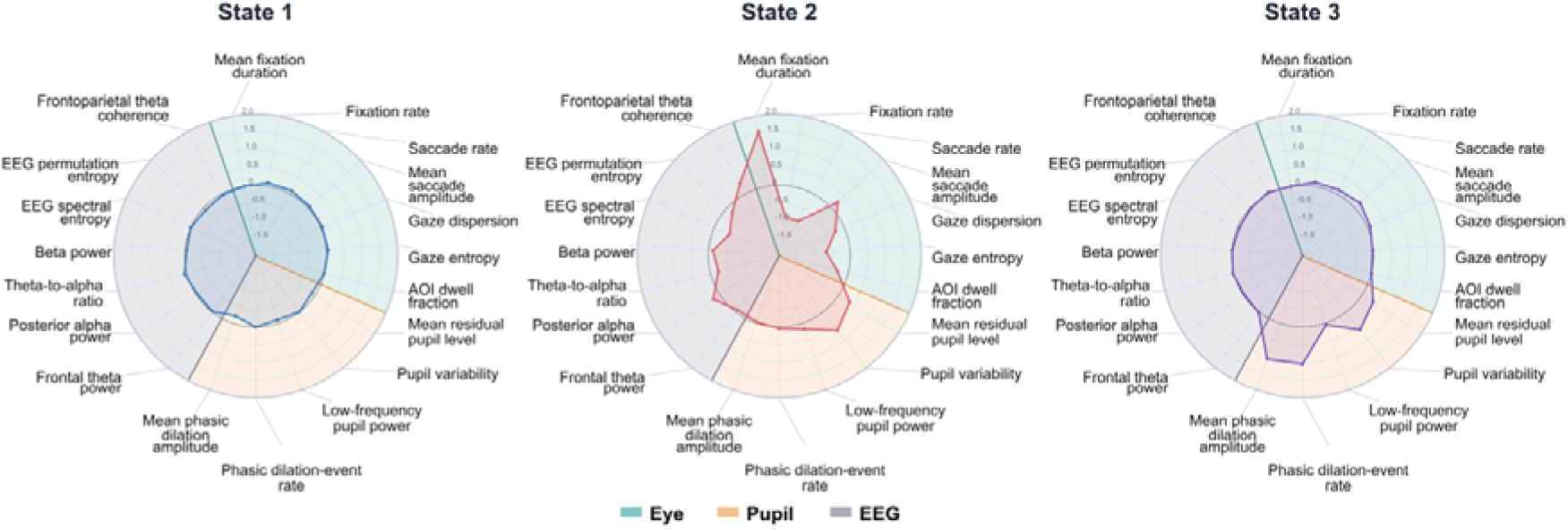
Multimodal profiles of the three latent states. Lines show participant-averaged, posterior-weighted means of the 19 standardized features on a common radial scale. The dashed circle marks zero, with larger radii indicating higher standardized feature values. Background sectors indicate eye-movement, pupil and EEG features. Low-frequency pupil power was computed over 0.05–0.20 Hz and log-transformed before standardization. EEG, electroencephalography; AOI, area of interest.

### 3.2 Temporal dynamics of multimodal states

State occupancy differed across states (Friedman χ²(2) = 108.097, *p* < .001), averaging 67.0%, 18.0%, and 15.0% for States 1, 2, and 3, respectively. State 1 had higher occupancy than State 2 (*t*(85) = 13.453, *p* < .001, *q* < .001) and State 3 (*t*(85) = 18.798, *p* < .001, *q* < .001); States 2 and 3 did not differ significantly (*t*(85) = 1.201, *p* = .233, *q* = .233; Figure 3A).

**Figure 3.**
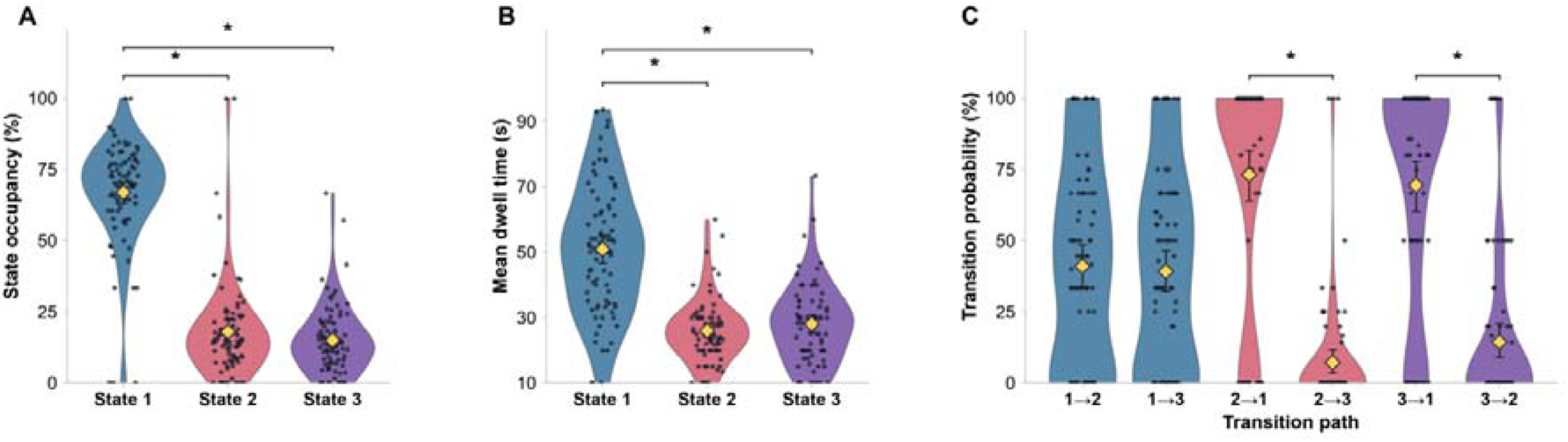
Temporal dynamics of the three latent states. Measures were derived from overlapping 30-s windows with a 10-s step. (A) Percentage of valid windows assigned to each state. (B) Mean dwell time, calculated from consecutive state runs. (C) Conditional transition probabilities excluding self-transitions; participants without a switch from the source state were assigned 0. Dots represent participants; violins show distributions; yellow diamonds indicate means, and error bars show participant-bootstrap 95% confidence intervals. Brackets indicate significant state comparisons in A and B and selected significant within-source destination comparisons in C (* *q* < .05). FDR correction was applied across the three pairwise comparisons separately for A and B and across all 15 pairwise comparisons for C; the complete set of transition comparisons is reported in Supplementary Table S2.

Mean dwell time also differed across states (Friedman χ²(2) = 79.908, *p* < .001). State 1 had longer dwell times than State 2 (*t*(73) = 12.432, *p* < .001, *q* < .001) and State 3 (*t*(74) = 10.704, *p* < .001, *q* < .001), whereas States 2 and 3 did not differ significantly (*t*(68) = -1.380, *p* = .172, *q* = .172; Figure 3B).

Zero-inclusive conditional transition probabilities differed across the six directed paths (Friedman χ²(5) = 181.216, *p* < .001). Across the 15 pairwise comparisons, 13 remained significant after Benjamini–Hochberg FDR correction. Notably, as shown in Figure 3C, within both States 2 and 3, transitions to State 1 were more probable than transitions to the alternative state (2→1 > 2→3: *t*(85) = 11.841, *p* < .001, *q* < .001; 3→1 > 3→2: *t*(85) = 8.471, *p* < .001, *q* < .001). In contrast, the two outgoing transitions from State 1 did not differ significantly (1→2 vs. 1→3: *t*(85) = 0.280, *p* = .780, *q* = .780). The complete set of 15 pairwise comparisons is reported in Supplementary Table S2.

### 3.3 Associations between relative state balance and quiz performance

In the adjusted participant-clustered GEE models, relative state balance of State 1 was not significantly associated with quiz performance (β = 0.036, *p* = .510, *q* = .510, 95% CI [-0.071, 0.143]; Figure 4A). In contrast, greater relative state balance of State 2 was associated with poorer quiz performance (β = -0.214, *p* < .001, *q* < .001, 95% CI [-0.330, -0.097]; Figure 4B). Greater relative state balance of State 3 was associated with better quiz performance (β = 0.151, *p* = .005, *q* = .008, 95% CI [0.045, 0.258]; Figure 4C).

**Figure 4.**
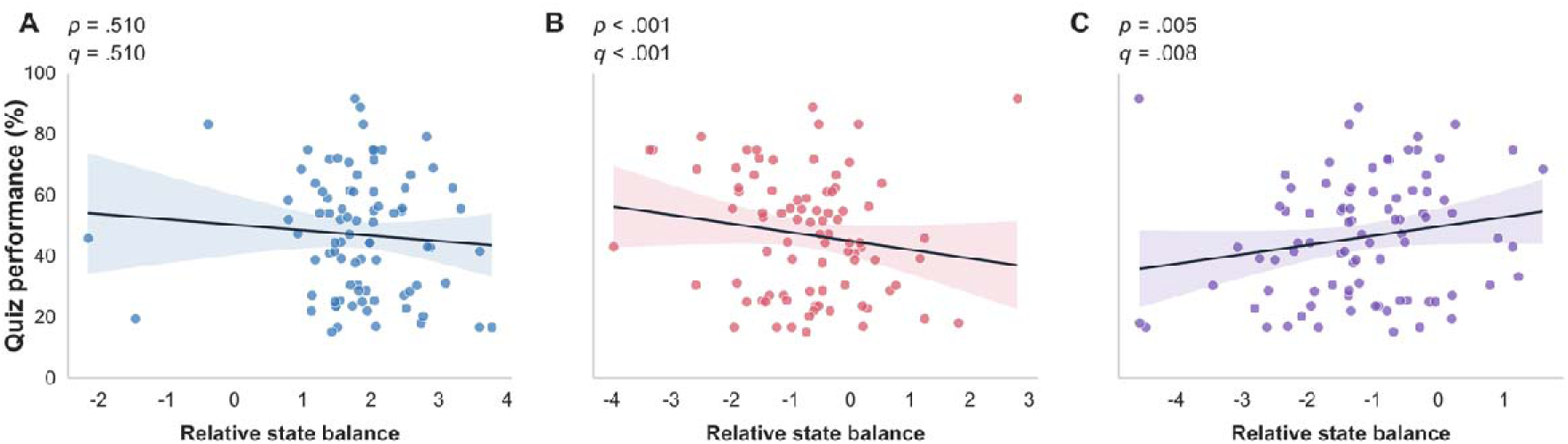
Associations between relative state balance and quiz performance for States 1 (A), 2 (B), and 3 (C), respectively. Relative state balance is the natural log of target-state posterior occupancy divided by the geometric mean of the other two occupancies, with 10 added to each occupancy, averaged across videos for each participant. Each dot represents a participant. Lines show unadjusted ordinary least-squares fits, with shaded pointwise 95% confidence intervals for the mean response. Annotations instead report two-sided *p* values from participant-clustered GEE models adjusted for age, sex and experiment-specific video identity, and corresponding *q* values after FDR correction across the three state-balance tests. The plotted lines are descriptive and are not fitted GEE predictions.

### 3.4 Associations between state transitions and quiz performance

Among the six directed between-state transition probabilities, only the probability of transitioning from State 3 to State 1 was positively associated with quiz performance after Benjamini–Hochberg FDR correction (β = 0.037, *p* = .001, *q* = .007, 95% CI [0.015, 0.060]). None of the other five transition probabilities remained significant after correction (*p* = .049–.653; *q* = .111–.653).

### 3.5 Window-length sensitivity analysis

Matched state profiles were similar across window lengths: State 1, *r* = .944; State 2, *r* = .976; and State 3, *r* = .991 (all nominal *p* < .001). The complete transition matrices, including self-transitions, were also highly correlated (*r* = .974, nominal *p* < .001).

In the 20-s common-record analysis, State 2 showed a negative state-balance association with quiz performance (β = -0.161, *p* = .009, *q* = .017, 95% CI [-0.283, -0.039]), whereas State 3 showed a positive association (β = 0.135, *p* = .011, *q* = .017, 95% CI [0.031, 0.239]). State 1 was not significantly associated with performance (β = -0.008, *p* = .899, *q* = .899, 95% CI [-0.136, 0.120]). Full estimates, including those from the common-record 30-s refit, are provided in Supplementary Table S3.

None of the six directed transition–quiz associations remained significant after FDR correction in the 20-s analysis (unadjusted *p* values = .022–.980; *q* values = .132–.980). Notably, the State 3 to State 1 association observed in the primary 30-s analysis was nominally significant in the 20-s analysis (*p* = .022) but did not remain significant after FDR correction (*q* = .132). Full results for both common-record refits are reported in Supplementary Table S4.

## 4 Discussion

The present findings indicate that learning during naturalistic video viewing is associated with the temporal organization of multimodal activity rather than with a single overall level of engagement. Three recurrent states with distinct combinations of eye-movement, pupil, and EEG features were identified. Relative state balance of State 2 and State 3 showed opposing associations with quiz performance, whereas relative state balance of State 1 was not associated with performance. In addition, the State 3-to-State 1 transition was positively associated with performance after correction. By distinguishing relative state balance from directed transition probabilities, the present analysis captures two complementary aspects of viewing dynamics: how activity is distributed across recurring configurations and how these configurations are linked over time. These findings extend previous work linking fluctuating neural states to cognitive performance (Esterman et al., 2013; Taghia et al., 2018; Vidaurre et al., 2016) and physiological engagement during educational videos to learning (Cohen et al., 2018; Madsen et al., 2021; Madsen & Parra, 2025), by examining their joint organization across ocular, pupillary, and neural signals.

State 3 showed a positive association between relative state balance and quiz performance and was distinguished primarily by more frequent and larger phasic pupil dilations. Phasic pupil responses have been linked to vigilance (Beatty, 1982) and transient engagement in adaptive-gain accounts (Aston-Jones & Cohen, 2005). Pupil dynamics have also been related to neuromodulatory activity (Reimer et al., 2016) and prioritized encoding (Clewett et al., 2018), although pupillometry does not provide a specific measure of locus-coeruleus activity (Joshi & Gold, 2020). Importantly, State 2 also showed elevated pupil level and variability but was associated with poorer performance, suggesting that pupil magnitude alone is unlikely to account for the observed behavioural pattern. Moreover, these state profiles describe standardized within-recording variation, not absolute arousal levels. State 3 may instead reflect transient changes in arousal, attentional engagement, or information updating that occur within the ongoing viewing process. One possibility is that appropriately timed episodes of phasic engagement are more relevant to learning than sustained physiological activation. Whether these episodes coincide with informative or otherwise salient events in the videos remains to be established.

The positive association between the State 3-to-State 1 transition and quiz performance provides a complementary perspective on this pattern. State 1 was the dominant background configuration, whereas State 3 was characterized primarily by phasic pupil activity. The transition may therefore capture a sequence in which a brief period of enhanced processing is followed by a return to the ongoing viewing configuration. This interpretation is broadly consistent with evidence that dynamic neural reconfiguration is related to cognitive performance (Shine et al., 2016; Taghia et al., 2018). Importantly, however, the transition probability used here is zero-inclusive: it reflects the observed destination of switching and assigns a value of zero when no switch from the source state occurs. It therefore does not directly measure switching frequency. The association should consequently be interpreted as evidence that the temporal context of a state may be relevant to learning, rather than as evidence that more frequent switching improves performance. More generally, the findings motivate further investigation of whether transient engagement followed by re-entry into an ongoing viewing state represents one aspect of adaptive temporal regulation during learning.

State 2 showed the opposite association with quiz performance and was characterized by longer fixations, lower fixation and saccade rates, lower gaze entropy, elevated pupil level and variability, and distinct EEG features including posterior alpha activity and frontoparietal theta coherence. The interpretation of these features is not straightforward. Gaze entropy is highly dependent on task and stimulus structure (Shiferaw et al., 2019), while alpha activity can reflect multiple aspects of attentional processing (Van Diepen et al., 2019). The joint profile may therefore indicate a mode of relatively sustained but constrained visual sampling, in which learners remain focused on a limited portion of the visual information as the content changes. Such stability may be adaptive under some task demands, and the present association should not be taken to indicate that reduced visual exploration is inherently detrimental. Rather, the finding suggests that the relationship between visual sampling and learning may depend on its broader physiological context. Importantly, the multimodal state should not be equated with disengagement, nor can the present data establish that any individual feature within the state impairs learning.

The 20-s and 30-s analyses showed similar state profiles and consistent opposing state-balance–performance associations. However, no transition–quiz association survived FDR correction in the common-record refits. These findings support the window-length stability of the state-balance associations but not the robustness of the transition–quiz finding. More broadly, the findings complement conventional average measures and previous studies linking physiological engagement to learning (Cohen et al., 2018; Madsen et al., 2021; Madsen & Parra, 2022, 2025) by providing a common temporal description of visual sampling, pupil activity, and neural dynamics. Removing stable recording-level differences and stimulus-shared responses before modelling further emphasized participant-specific variation around the response common across viewers. Thus, rather than asking whether a learner was generally more or less engaged, this approach asked how multiple physiological signals were organized over time during viewing. Whether such multimodal configurations capture information about learning that is not available from individual features or conventional summary measures remains an important question for future work and requires direct comparison with simple models.

Several limitations should be considered when interpreting these findings. First, the associations between multimodal states and quiz performance are observational. Prior knowledge, motivation, task engagement, and other unmeasured differences could contribute to both state organization and subsequent performance. Second, HMM states are statistical configurations that depend on feature selection, preprocessing, and model assumptions; overlapping windows also limit temporal precision and may obscure rapid event transitions. Finally, the study focused on short educational videos and quiz performance assessed after viewing, so the findings may not generalize to longer learning episodes or indicate durable knowledge retention. Future studies should examine longer learning materials and include delayed assessments to test the generalizability and persistence of these associations.

In conclusion, this study identified three recurrent multimodal states in eye-movement, pupil, and EEG recordings during educational video viewing. Their associations with subsequent quiz performance differed: greater relative state balance of State 3, characterized primarily by more frequent and larger phasic pupil dilations, was associated with better performance, whereas greater relative state balance of State 2, characterized by more constrained visual sampling and a distinct neural profile, was associated with poorer performance. State 1 was the dominant configuration, but showed no detected association with performance. Among the directed transitions, only the State 3-to-State 1 transition was positively associated with performance after correction, suggesting that the temporal context of multimodal configurations may provide information complementary to the analysis of relative state balance. State profiles and state-balance associations were consistent across 20-s and 30-s windows, whereas the transition–quiz association was not retained after correction in common-record refits and requires replication. Together, these findings suggest that learning during naturalistic video viewing is associated with the relative expression and temporal organization of multiple ocular, pupillary, and neural signals rather than with a single sustained level of engagement. Transient engagement followed by a return to ongoing viewing provides one plausible interpretation of the observed pattern, but remains a hypothesis for future investigation. Event-resolved and experimentally controlled studies are needed to determine whether such temporal organization plays a functional role in learning.

## Acknowledgments

This study was supported by the Guangdong Basic and Applied Basic Research Foundation (2026A1515011529) awarded to Y.X.

## Author contributions: CRediT

**Ziqiang Tian**: Data curation, Formal analysis, Methodology, Software, Visualization, Writing - original draft, Writing - review & editing. **An Long**: Writing - original draft, Writing - review & editing. **Yaqiong Xiao**: Conceptualization, Funding acquisition, Project administration, Supervision, Writing - original draft, Writing - review & editing.

## Data and code availability statement

The preprocessed data used in this study are publicly available at https://doi.org/10.15387/fcp_indi.retro.bbbd. The analysis code used in this study has now been deposited in a public repository (https://github.com/Yaqiongxiao/BBBD.HMM).

