## Supplementary Materials for "Multimodal dynamics during naturalistic learning are associated with subsequent performance"

**Table S1. Detailed feature estimates and state comparisons**

| **Feature** | **State 1** | **State 2** | **State 3** | **Omnibus *p* / *q*** | **Pairwise comparisons (*p* / *q*)** | | |
| --- | --- | --- | --- | --- | --- | --- | --- |
|  |  |  |  |  | **State 1 vs. 2** | **State 1 vs. 3** | **State 2 vs. 3** |
| Mean fixation duration | -0.001 ± 0.235 | 1.555 ± 1.057 | -0.007 ± 0.486 | *p* < .001 *q* < .001 | *p* < .001 *q* < .001 | *p* = .866 *q* = .903 | *p* < .001 *q* < .001 |
| Fixation rate | 0.098 ± 0.157 | -0.900 ± 0.560 | 0.119 ± 0.421 | *p* < .001 *q* < .001 | *p* < .001 *q* < .001 | *p* = .604 *q* = .717 | *p* < .001 *q* < .001 |
| Saccade rate | 0.105 ± 0.150 | -0.886 ± 0.548 | 0.126 ± 0.431 | *p* < .001 *q* < .001 | *p* < .001 *q* < .001 | *p* = .591 *q* = .717 | *p* < .001 *q* < .001 |
| Mean saccade amplitude | 0.063 ± 0.188 | 0.239 ± 0.656 | 0.207 ± 0.543 | *p* = .219 *q* = .245 | *p* = .030 *q* = .052 | *p* = .021 *q* = .038 | *p* = .870 *q* = .903 |
| Gaze dispersion | 0.044 ± 0.148 | -0.277 ± 0.691 | 0.082 ± 0.432 | *p* = .007 *q* = .010 | *p* < .001 *q* = .001 | *p* = .871 *q* = .903 | *p* < .001 *q* < .001 |
| Gaze entropy | 0.047 ± 0.191 | -0.669 ± 0.632 | -0.020 ± 0.420 | *p* < .001 *q* < .001 | *p* < .001 *q* < .001 | *p* = .337 *q* = .458 | *p* < .001 *q* < .001 |
| AOI dwell fraction | -0.009 ± 0.196 | -0.315 ± 0.740 | 0.004 ± 0.341 | *p* = .027 *q* = .036 | *p* = .004 *q* = .008 | *p* = .636 *q* = .740 | *p* = .001 *q* = .003 |
| Mean residual pupil level | -0.105 ± 0.147 | 0.372 ± 0.531 | 0.373 ± 0.821 | *p* < .001 *q* < .001 | *p* < .001 *q* < .001 | *p* < .001 *q* < .001 | *p* = .991 *q* = .991 |
| Pupil variability | -0.002 ± 0.193 | 0.666 ± 0.567 | 0.639 ± 0.785 | *p* < .001 *q* < .001 | *p* < .001 *q* < .001 | *p* < .001 *q* < .001 | *p* = .576 *q* = .714 |
| Low-frequency pupil power | -0.100 ± 0.182 | 0.181 ± 0.439 | 0.049 ± 0.554 | *p* < .001 *q* < .001 | *p* < .001 *q* < .001 | *p* = .008 *q* = .016 | *p* = .271 *q* = .396 |
| Phasic dilation-event rate | 0.001 ± 0.181 | 0.047 ± 0.385 | 1.050 ± 0.802 | *p* < .001 *q* < .001 | *p* = .314 *q* = .437 | *p* < .001 *q* < .001 | *p* < .001 *q* < .001 |
| Mean phasic dilation amplitude | -0.215 ± 0.211 | -0.037 ± 0.535 | 1.078 ± 1.835 | *p* < .001 *q* < .001 | *p* = .009 *q* = .017 | *p* < .001 *q* < .001 | *p* < .001 *q* < .001 |
| Frontal theta power | -0.018 ± 0.231 | -0.075 ± 0.865 | 0.017 ± 0.389 | *p* = .442 *q* = .466 | *p* = .561 *q* = .710 | *p* = .825 *q* = .903 | *p* = .728 *q* = .830 |
| Posterior alpha power | -0.032 ± 0.182 | 0.222 ± 0.634 | -0.026 ± 0.328 | *p* = .066 *q* = .084 | *p* = .006 *q* = .012 | *p* = .746 *q* = .834 | *p* = .026 *q* = .046 |
| Theta-to-alpha ratio | 0.071 ± 0.209 | -0.230 ± 0.629 | 0.026 ± 0.329 | *p* < .001 *q* = .001 | *p* < .001 *q* = .001 | *p* = .469 *q* = .607 | *p* = .007 *q* = .015 |
| Beta power | -0.046 ± 0.185 | -0.122 ± 0.605 | -0.009 ± 0.450 | *p* = .142 *q* = .169 | *p* = .174 *q* = .267 | *p* = .991 *q* = .991 | *p* = .309 *q* = .437 |
| EEG spectral entropy | 0.001 ± 0.194 | -0.472 ± 0.723 | -0.046 ± 0.362 | *p* < .001 *q* < .001 | *p* < .001 *q* < .001 | *p* = .177 *q* = .267 | *p* < .001 *q* < .001 |
| EEG permutation entropy | -0.084 ± 0.252 | -0.197 ± 0.547 | -0.018 ± 0.383 | *p* = .522 *q* = .522 | *p* = .178 *q* = .267 | *p* = .399 *q* = .530 | *p* = .041 *q* = .069 |
| Frontoparietal theta coherence | -0.040 ± 0.159 | 0.326 ± 0.535 | 0.043 ± 0.298 | *p* < .001 *q* < .001 | *p* < .001 *q* < .001 | *p* = .121 *q* = .197 | *p* = .001 *q* = .003 |

Note. State values are M ± SD of eligible participant-level posterior-weighted standardized feature estimates; SD is the sample standard deviation. Low-frequency pupil power is log-transformed power over 0.05–0.20 Hz. Profiles describe relative within-recording variation.

**Table S2. Comparisons among directed transition probabilities**

| **Paths** | **Mean difference** | ***t*** | **df** | ***p*** | ***q*** | **95% CI** |
| --- | --- | --- | --- | --- | --- | --- |
| 1→2 vs. 1→3 | 0.018 | 0.280 | 85 | .780 | .780 | [-0.109, 0.144] |
| 1→2 vs. 2→1 | -0.321 | -7.095 | 85 | < .001 | < .001 | [-0.411, -0.231] |
| 1→2 vs. 2→3 | 0.339 | 7.279 | 85 | < .001 | < .001 | [0.246, 0.431] |
| 1→2 vs. 3→1 | -0.283 | -4.564 | 85 | < .001 | < .001 | [-0.407, -0.160] |
| 1→2 vs. 3→2 | 0.266 | 5.909 | 85 | < .001 | < .001 | [0.177, 0.356] |
| 1→3 vs. 2→1 | -0.339 | -6.991 | 85 | < .001 | < .001 | [-0.435, -0.242] |
| 1→3 vs. 2→3 | 0.321 | 7.258 | 85 | < .001 | < .001 | [0.233, 0.409] |
| 1→3 vs. 3→1 | -0.301 | -6.691 | 85 | < .001 | < .001 | [-0.391, -0.212] |
| 1→3 vs. 3→2 | 0.248 | 5.088 | 85 | < .001 | < .001 | [0.151, 0.346] |
| 2→1 vs. 2→3 | 0.660 | 11.841 | 85 | < .001 | < .001 | [0.549, 0.771] |
| 2→1 vs. 3→1 | 0.038 | 0.711 | 85 | .479 | .513 | [-0.068, 0.143] |
| 2→1 vs. 3→2 | 0.587 | 11.336 | 85 | < .001 | < .001 | [0.484, 0.690] |
| 2→3 vs. 3→1 | -0.622 | -12.541 | 85 | < .001 | < .001 | [-0.721, -0.524] |
| 2→3 vs. 3→2 | -0.073 | -2.166 | 85 | .033 | .038 | [-0.139, -0.006] |
| 3→1 vs. 3→2 | 0.550 | 8.471 | 85 | < .001 | < .001 | [0.421, 0.679] |

**Table S3. Associations between relative state balance and quiz performance in common-record** **20-s and 30-s windows**

| **Window (s)** | **State** | **β** | ***p*** | ***q*** | **95% CI** |
| --- | --- | --- | --- | --- | --- |
| 20 | 1 | -0.008 | .899 | .899 | [-0.136, 0.120] |
| 20 | 2 | -0.161 | .009 | .017 | [-0.283, -0.039] |
| 20 | 3 | 0.135 | .011 | .017 | [0.031, 0.239] |
| 30 | 1 | -0.005 | .940 | .940 | [-0.123, 0.114] |
| 30 | 2 | -0.231 | < .001 | < .001 | [-0.354, -0.108] |
| 30 | 3 | 0.166 | .001 | .002 | [0.064, 0.268] |

**Table S4. Associations between state transitions and quiz performance in common-record** **20-s and 30-s windows.**

| **Window (s)** | **Path** | **β** | ***p*** | ***q*** | **95% CI** |
| --- | --- | --- | --- | --- | --- |
| 20 | 1→2 | -0.003 | .829 | .980 | [-0.033, 0.026] |
| 20 | 1→3 | 0.013 | .313 | .480 | [-0.012, 0.038] |
| 20 | 2→1 | -0.0005 | .980 | .980 | [-0.037, 0.036] |
| 20 | 2→3 | 0.015 | .320 | .480 | [-0.015, 0.045] |
| 20 | 3→1 | 0.033 | .022 | .132 | [0.005, 0.062] |
| 20 | 3→2 | -0.014 | .269 | .480 | [-0.038, 0.011] |
| 30 | 1→2 | -0.025 | .101 | .304 | [-0.055, 0.005] |
| 30 | 1→3 | 0.027 | .072 | .304 | [-0.002, 0.057] |
| 30 | 2→1 | -0.004 | .876 | .964 | [-0.048, 0.041] |
| 30 | 2→3 | -0.001 | .964 | .964 | [-0.034, 0.032] |
| 30 | 3→1 | 0.017 | .181 | .362 | [-0.008, 0.043] |
| 30 | 3→2 | -0.008 | .516 | .774 | [-0.030, 0.015] |
